# Information theoretic analysis of hyperspectral imaging systems with applications to fluorescence microscopy

**DOI:** 10.1101/373894

**Authors:** Sripad Ram

**Affiliations:** Global Pathology, Drug Safety Research and Development, Pfizer Inc., 10646 Science Center Drive, San Diego, CA 92121 USA

**Keywords:** Spectral unmixing, Multispectral imaging, Multiplex imaging, Cramer-Rao lower bound

## Abstract

We present a general stochastic model for hyperspectral imaging data and derive analytical expressions for the Fisher information matrix for the underlying spectral unmixing problem. We investigate the linear mixing model as a special case and define a linear unmixing performance bound by using the Cramer-Rao inequality. As an application, we consider fluorescence imaging and show how the performance bound provides a spectral resolution limit that predicts how accurately a pair of spectrally similar fluorescent labels can be spectrally unmixed. We also report a novel result that shows how the spectral resolution limit can be overcome by exploiting the phenomenon of anti-Stokes shift fluorescence. In addition, we investigate how photon statistics, channel addition and channel splitting affect the performance bound. Finally by using the performance bound as a benchmark, we compare the performance of the least squares and the maximum likelihood estimators for spectral unmixing. For the imaging conditions tested here, our analysis shows that both estimators are unbiased and that the standard deviation of the maximum likelihood estimator is consistently closer to the performance bound than that of the least squares estimator. The results presented here are based on broad assumptions regarding the underlying data model and are applicable to hyperspectral data acquired with point detectors, sCMOS, CCD and EMCCD imaging detectors.

**EDICS:** ELI-COL, COI-MCI.

## I. Introduction

Hyperspectral imaging represents a broad class of techniques that capture spectral and spatial data from the object of interest. Fluorescence microscopy is a powerful tool to study microscopic objects such as biological cells at high spatial and temporal resolution. Fluorescence imaging supports the visualization of multiple targets within the object of interest, which is facilitated by labeling the targets with fluorescent labels that have distinct excitation and emission spectra (Fig. 1A). Consequently, the need for hyperspectral imaging arises in such applications when the emission spectra of the fluorescent labels significantly overlap (Fig. 1B). There exist a variety of approaches to carry out hyperspectral imaging in a fluorescence microscope. For instance, hyperspectral imaging can be carried out either with a confocal ([1], [2], [3]) or a linescanning ([4]) microscope that has spectral detection capability. Alternately, it can also be carried out on a widefield microscope by using either narrowband excitation and emission filters ([5], [6], [7], [8]) or by using an electronically controlled liquid crystal tunable filter ([9], [10]). Common to all these techniques is the underlying hyperspectral data which consists of a sequence of 2D images acquired at different spectral windows. Thus given the hyperspectral data, the goal is then to estimate the relative abundance (typically represented in photon counts) of the different labels at each pixel, and this is referred to as the spectral unmixing problem (Fig. 1C).

**Fig. 1.**
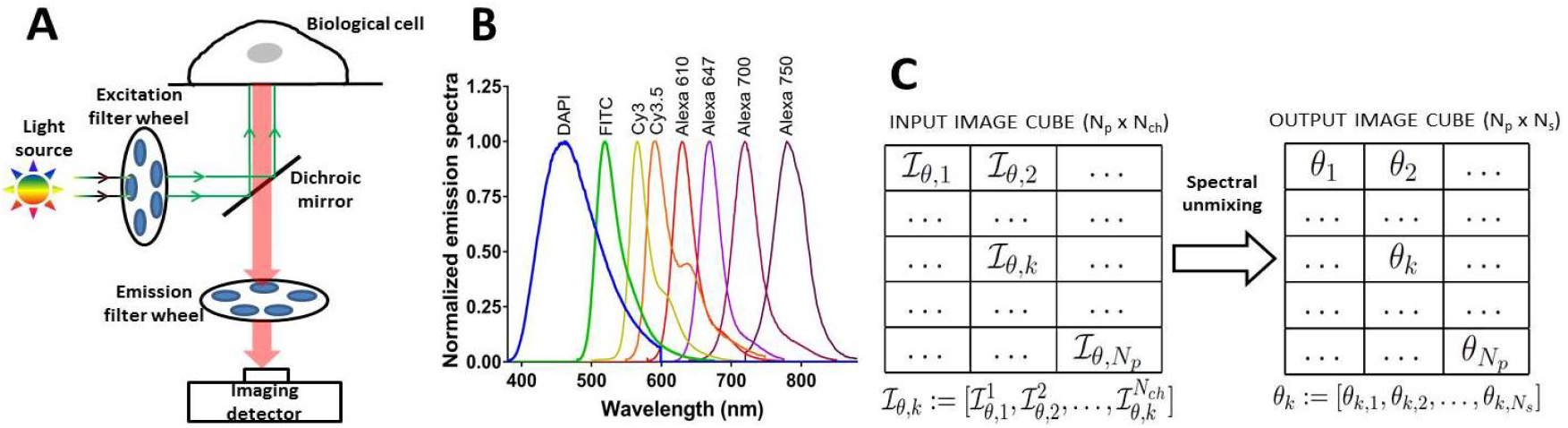
Hyperspectral imaging in fluorescence microscopy. Panel A shows a typical configuration of a fluorescence microscope for multicolor imaging application. The sample is selectively illuminated at distinct excitation passbands (excitation filter wheel) and the fluorescence signal is then collected at matched emission passbands (emission filter wheel) on an imaging detector. Panel B shows the emission spectra of fluorescent labels exhibiting significant spectral overlap. Panel C illustrates the spectral unmixing problem where given a hyperspectral data set, i.e. the input image cube, which consists of *N_ch_* spectral images each with *N_p_* pixels, the goal is to obtain an estimate of the output image cube that represents the relative abundance *θ_k_* of *N_s_* different fluorescent labels at each pixel in the sample. Here ℐ_*θ,k*_ is a random vector with probability density *p_θ,k_* that is a function of *ν_θ,k_*, which describes the signal at the *k^th^* pixel in the input image cube.

A common approach to solving the spectral unmixing problem assumes that the number of labels present in the sample is known. Spectral unmixing approaches that make use of this strategy include the least squares estimator ([11], [12], [13]), the maximum likelihood estimator ([14], [15]), and phasor based approaches ([3], [16]). In fluorescence imaging applications, we typically assume a linear mixing model i.e., *μ_θ,k_* = *Aθ_k_*, where *μ_θ,k_*, which describes the signal detected at the k^*th*^ pixel in the input image cube (see Fig. 1C), is a linear combination of the relative abundance *θ_k_* of the different fluorescent labels present at that pixel in the sample. Here A denotes the mixing matrix that specifies the relative contribution of each fluorescent label in every spectral channel. The mixing matrix depends on the optical configuration of the hypserspectal microscope and the spectral properties of the fluorescent dye. The mixing matrix can be computed either theoretically or experimentally by performing a calibration experiment with single color control samples.

A fundamental question that arises in hyperspectral imaging applications concerns with its performance limits which deals with the best possible accuracy with which the relative abundance of the different labels can be determined. Knowledge of the performance limit is important as it provides a metric to design and optimize a hyperspectral imaging system. Moreover, it also acts a benchmark to compare the performance of different spectral unmixing algorithms for a given hyperspectral dataset. In this paper, we present results to compute the performance limits of a hyperspectral imaging system. We adopt a stochastic framework to model the acquired data and derive the Fisher information matrix for the spectral unmixing problem. We then consider the linear mixing model as a special case. Using the Cramer-Rao inequality ([17]), we define a linear unmixing performance bound which provides a lower bound to the accuracy with which the relative abundance of the different labels can be estimated.

In the past, several groups have investigated the spectral unmixing problem. In [18], [19], the authors calculated the Fisher information matrix for a deterministic data model that is corrupted by white noise (Gaussian data model). In [15], [20], the authors derived the Fisher information matrix for a data model that is corrupted by shot noise (Poisson data model). The present manuscript provides a broad stochastic framework that is applicable to several different data models that are typically used to describe data from an imaging detector. Specifically, in addition to the above data models our results are also applicable for the Poisson + Gaussian data model and for the stochastic Poisson + Gaussian data model which are typically used to describe imaging data acquired from sCMOS/CCD cameras and EMCCD cameras, respectively.

The paper is organized as follow. In Section II we present the problem formulation, introduce the different data models, and derive the general expression of the Fisher information matrix for the spectral unmixing problem. We then investigate conditions under which the Fisher information matrix is block diagonal as block diagonality has several implications. Next, we introduce the linear mixing model and derive the Fisher information matrix for the same. We also derive analytical expressions of the Fisher information matrix that pertain to the different data models. Further we derive results to address questions regarding the design of hyperspectral imaging systems such as the addition or splitting of spectral channels and its impact on spectral unmixing.

In Section IV, we illustrate the results derived in Section II by considering different hyperspectral imaging configurations and fluorescent label pairs. We show that for a pair of regular fluorescent labels there exists is a spectral resolution limit, which predicts how spectrally close the emission spectra of the two labels can be and still be accurately spectrally resolved. We also show that by exploiting the phenomenon of anti-Stokes shift fluorescence the spectral resolution limit can be overcome and that the emission spectra of the fluorescent labels can be arbitrarily close to each other. We also illustrate how photon statistics, and the addition or splitting of spectral channels can affect the performance bound for different data models. In Section V, we compare the performance of two spectral unmixing algorithms, namely the least squares and maximum likelihood estimators, on hyperspectral data. Our results show that for the imaging conditions tested here, both estimators are unbiased and their performance come close to the theoretical performance bound with the maximum likelihood estimator being consistently closer to the performance bound than the least squares estimator.

## II. Theory

### A. Problem formulation

We consider a generic model of a hyperspectral imaging system wherein the sample is illuminated by a light source and the light that is transmitted or emitted by the sample is collected into spectral channels and recorded as a sequence of 2D spectral images, which we refer to as the input image cube. We assume that the size of every spectral image is the same. Let *N_p_* denote the total number of pixels in a spectral image and *N_ch_* denote the number of spectral images. We define a pixel block as a sequence of *N_ch_* pixels with the same pixel index, say *k* for *k* = 1, …, *N_p_*, across *N_ch_* different spectral images. We assume that the object of interest contains *N_s_* distinct but spectrally overlapping labels. Due to the presence of multiple labels, the signal at the *k^th^* pixel block can be described as a superposition of the relative abundance or photons detected from the individual labels that are present at that pixel in the sample. The spectral unmixing problem can then be stated as follows: given an input image cube which is a *N_p_* × *N_ch_* dimensional dataset, the goal is to estimate an output (or spectrally unmixed) image cube *θ* which is a *N_p_* × *N_s_* dimensional dataset that contains to the photon counts of *N_s_* labels at each pixel in the sample.

### B. Image formation model

Let 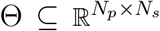 be the parameter space and *θ* ∈ Θ be the unknown parameter vector. The signal detected in the input image cube is modeled as a sequence of random vectors given by

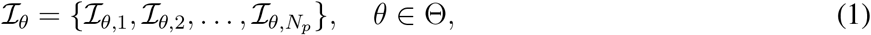

where 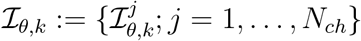 for *k* = 1, …, *N_p_*, and 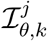 denotes an independent random variable with probability density function 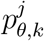 that models the detected photons at the *k^th^* pixel in the *j^th^* spectral image. Here we assume that for *θ ∈* Θ, *j* = 1, …, *N_ch_* and *k* = 1, …, *N_p_*,

**A1** 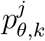 satisfies the regularity conditions ([17]) required for the calculation of the Fisher information matrix, and

**A2** 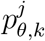 depends on *θ* through a non-negative function 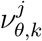 that is differentiable with respect to *θ*, i.e., 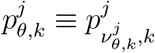.

As we will see, the term 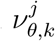 will typically describe the expected number of detected photons at the *k^th^* pixel in the *j^th^* spectral image for θ ∈ Θ, *j* = 1, …, *N_ch_* and *k* = 1, …, *N_p_*. We note that 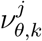 in turn can be expressed as a function of *θ* which we wish to estimate.

### C. Specific data models

We note that the above assumptions regarding the probability density function 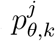 are quite broad. To illustrate how this covers a wide variety of imaging conditions, we consider the following data models.

#### Gaussian data model

Here we consider the case where the photons detected at the *k^th^* pixel in the *j^th^* spectral image is modeled as a deterministic signal 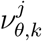 that is corrupted by additive Gaussian noise. Hence 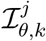 is given by 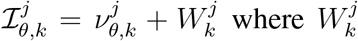 is an independent Gaussian random variable with mean 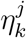 and standard deviation 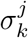 for *j* = 1, …, *N_ch_* and *k* = 1, …, *N_p_*. Then the probability density function of 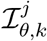 is given by

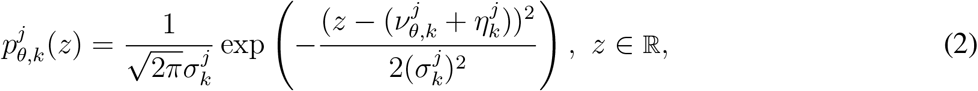

*θ* ∈ Θ, *j* = 1, …, *N_ch_*, *k* = 1, …, *N_p_*. For the above equation, it is straightforward to verify that assumptions **A1** and **A2** are satisfied.

#### Poisson data model

Here we assume that the detected photons at each pixel are Poisson distributed. Thus we have 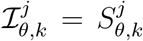, where 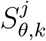 is an independent Poisson random variable with mean 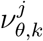 for *θ* ∈ Θ, *j* = 1, …, *N_ch_* and *k* = 1, …, *N_p_*. he probability density function of 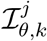 is given by

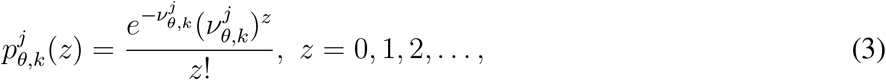

*θ* ∈ Θ, *j* = 1, …, *N_ch_ k* = 1, …, *N_p_*. It immediately follows that assumptions **A1** and **A2** are satisfied. For instance, this specific data model would be applicable for imaging systems such as a confocal or a multiphoton microscope, where the main source of randomness in the data is attributed to shot noise statistics ([21]).

#### Poisson + Gaussian data model

Here we consider 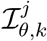 to be given by 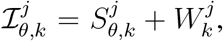 for *θ* ∈ Θ *j* = 1, …, *N_ch_*, and *k* = 1, …, *N_p_*. Here, 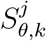 is Poisson random variable with mean 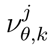 that models the detected photons at the *k^th^* pixel in the *j^th^* spectral image, and 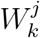 is an independent Gaussian random variable with mean 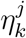 and standard deviation 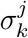 that models the measurement noise at the *k^th^* pixel in the *j^th^* spectral image, for *θ* ∈ Θ, *j* = 1, …, *N_ch_* and *k* = 1, …, *N_p_*. Here we assume that 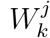 is independent of *θ* for *θ* ∈ Θ, *j* = 1, …, *N_ch_* and *k* = 1, …, *N_p_*. Then the probability density function of 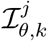 is given by ([22])

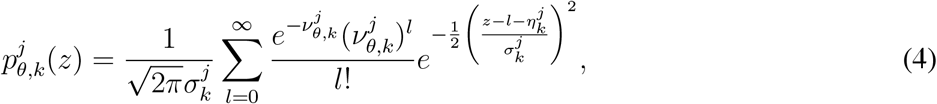

for *z* ∈ ℝ, *θ* ∈ Θ, *k* = 1, …, *N_p_* and *j* = 1, …, *N_ch_*. From the above expression it immediately follows that assumption **A2** is satisfied. Moreover, it can also be shown that *p_θ_* satisfies the regularity conditions ([23]) thereby satisfying assumption **A1**. This data model is applicable to a widefield microscope configuration in which the images are acquired with either a CCD or a sCMOS camera. Here, in addition to the shot noise statistics the detected signal is also corrupted by the measurement noise of the detector.

#### Stochastic-Poisson + Gaussian data model

For completeness, we also consider another data model where in addition to shot noise statistics and measurement noise of the detector, the model takes into account stochastic signal amplification, which, for example, occurs in an electron multiplying CCD camera. Specifically, the detected photons at each pixel that is Poisson distributed with mean 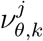 is amplified by a random function **M** that is independent of *θ* and the amplified signal is further corrupted by additive Gaussian noise ([24]), for *θ* ∈ Θ, *j* = 1, …, *N_ch_* and *k* = 1, …, *N_p_* . **M** is typically modeled as a branching process ([25]) with the initial particle count to be Poisson distributed (with mean 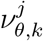) and the individual offspring count to be a zero modified geometric distribution. For the above data model, the probability density function can be written as (see [24] for details)

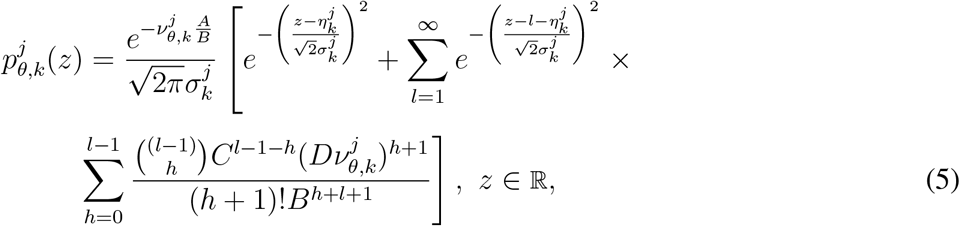

where *A*, *B*, *C* and *D* are constants that are independent of *θ* for *θ* ∈ Θ, *j* = 1, …, *N_ch_* and *k* = 1, …, *N_p_*. From the above equation we see that assumption **A2** is satisfied. Further, it can be shown that the above equation satisfies assumption **A1** ([24]).

### D. General expression for the Fisher information matrix

In this section, we derive a general expression of the Fisher information matrix for the output image cube *θ* using the image formation model described in Section II-B. We also investigate conditions under which the Fisher information matrix **I**(*θ*) is diagonal, as diagonality of **I**(*θ*) has several implications. In the present context, since *θ* is a vector with *N_p_* × *N_s_* elements, the resulting Fisher information matrix will be a (*N_p_* × *N_s_*) *×* (*N_p_* × *N_s_*) matrix, which can be very large. Thus a diagonal **I**(*θ*) can render its calculation to be more tractable. In general for an n-dimensional parameter vector *θ* = (*θ*_1_, …, *θ_n_*) ∈ Θ, if I(*θ*) is diagonal, then the limit of the accuracy of *θ_i_* is independent of other components of *θ*.

Here, the parameter vector is given by

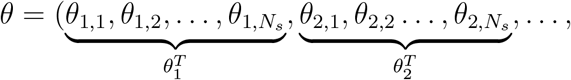

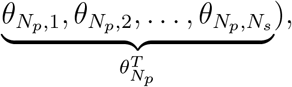

where *θ_k_* = (*θ_k,_*_1_, *θ_k,_*_2_, …, *θ_k,N_*)^*T*^ denotes the unknown photon counts of the *N_s_* different labels at the *k^th^* pixel in the output image cube for *k* = 1 …, *N_p_*. Further we assume that for *j* = 1, …, *N_ch_* and *k* ≠ *m* and 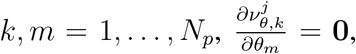 *θ* ∈ Θ where 0 denotes a 1 × *N_s_* row vector with all elements equal to zero. The validity of this assumption becomes evident in the next section where we will consider an explicit relationship between *ν_θ,k_* and the components of *θ_k_* for the linear mixing model. As we will see, the above assumption results in the Fisher information matrix being block diagonal, where the diagonal entries pertain to the Fisher information matrix of *θ_k_* for *k* = 1, …, *N_p_*.

In the following theorem, we state two results. The first result is a general expression for the Fisher information matrix pertaining to the image cube ℐ *_θ_*, which is analogous to a previously published result [24], [26]. The second result investigates conditions for block diagonality of the Fisher information matrix.

#### Theorem 2.1

Let 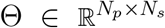 denote the parameter space. For *θ* ∈ Θ, let 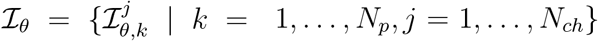 denote the input image cube that is defined in eq. 1. Assume that conditions **A1 - A2** are satisfied (see Section II-B).

1. For *θ* ∈ Θ, the Fisher information matrix for the output image cube *θ* is given by

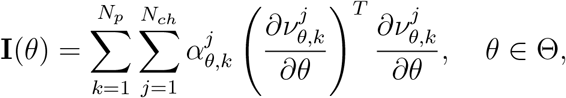

where for *θ* ∈ Θ *j* = 1,…, *N*_ch_ and *k* = 1,…, *N*_p_

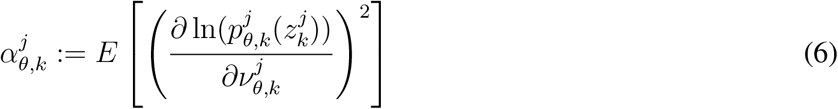

and 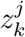 denotes a realization of 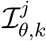.
2. Assume that for *k ≠m*, *k, m* = 1, …, *N_p_* and 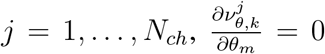, *θ* ∈ Θ. Then the Fisher information matrix given in result 1 of this Theorem can be written as

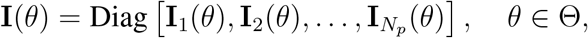

where for *θ* ∈ Θ and *k* = 1, …, *N_p_*

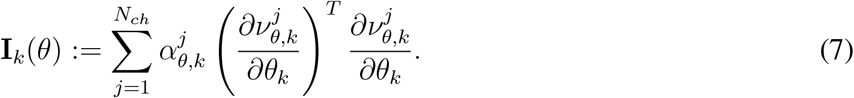

**Proof:**

1. See [24], [26] for proof.
2. For *θ* ∈ Θ and *k* = 1, …, *N_p_*, define

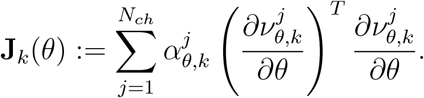

Then the Fisher information matrix given in result 1 of this Theorem can be written as

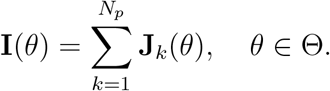

For *θ* ∈ Θ, *k* = 1, …, *N_p_* and *j* = 1, …, *N_ch_*, consider the term

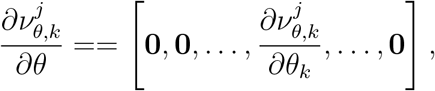

where we have used the fact that for 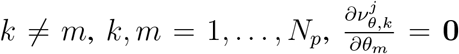 for *θ* ∈ Θ and 0 denotes a 1 × *N_s_* vector with all elements equal to zero. Using the above result, we then have

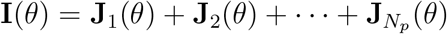

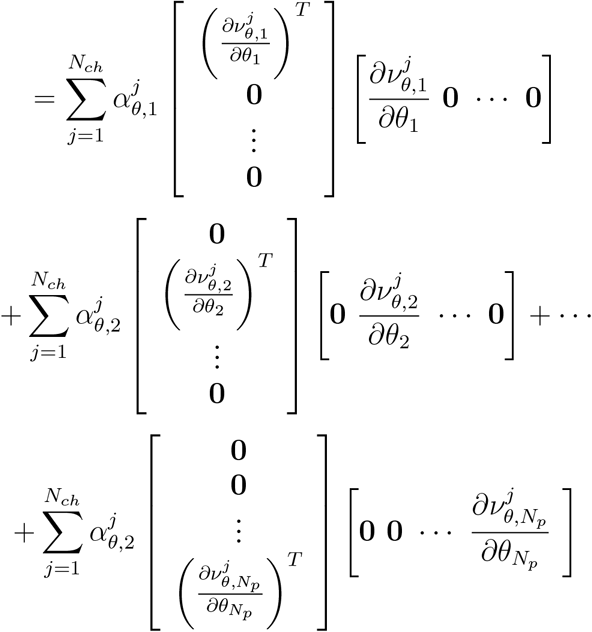

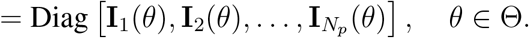

From result 2 of the above theorem we see that the block diagonal representation reduces the computational complexity to calculate the Fisher information matrix. Specifically, for *k* = 1, …, *N_p_* the *k^th^* block matrix **I**_*k*_(*θ*) is a *N_s_* × *N_s_* square matrix, which is relatively straightforward to compute.

### E. Linear mixing model and the mixing matrix

In the previous theorem we derived a general expression of the Fisher information matrix for the output image cube *θ* pertaining to a general hyperspectral imaging system. Here we next consider the case when the non negative function 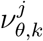 is a linear superposition of the components of the unknown parameter vector *θ* for *j* = 1, …, *N_ch_* and *k* = 1, …, *N_p_*. For *θ* ∈ Θ, define 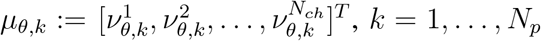, *k*=1,…, *N_p_*. Then for the linear mixing model we have

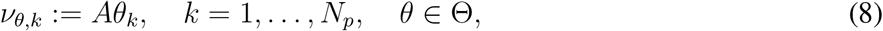

where A is a *N_ch_* × *N_s_* dimensional matrix with elements {*a_ij_*; *i* = 1, …, *N_ch_*; *j* = 1, …, *N_s_*} known as the *mixing matrix* and *θ_k_* = [*θ_k,_*_1_*, θ_k,_*_2_, …, *θ_k,N_*]^*T*^, *k* = 1, …, *N_p_*. The columns of A pertain to the different labels, the rows of A pertain to the different spectral channels, and the element *a_ij_* denotes the contribution of the *j^th^* label in the *i^th^* channel. As we will see, the mixing matrix will play an important role is governng the behavior of the Fisher information matrix for the linear mixing model. We note that the linear mixing model is used to model hyperspectral data in many applications including fluorescence microscopy.

In the following Theorem we derive the Fisher information matrix for the linear mixing model. We also consider a special case where *ν_θ,k_* = *θ_k_* for *θ* ∈Θ, *k* = 1, …, *N_p_* and *N_ch_* = *N_s_*. This special case pertains to an ideal imaging configuration where the signal from a given label can be collected without being corrupted by signal from other labels at every pixel block in the input mage cube.

#### Theorem 2.2

Let **I**(*θ*) denote the Fisher information matrix of the output image cube *θ* that is given in Theorem 2.1.

1. For *θ* ∈ Θ and *k* = 1, …, *N_p_* let *ν_θ,k_* be given by eq. 8. Then the Fisher information matrix of the output image cube *θ* for the linear mixing model is given by

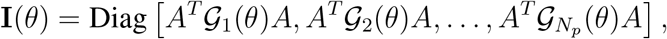

where *θ* ∈ Θ, *A* denotes the mixing matrix,

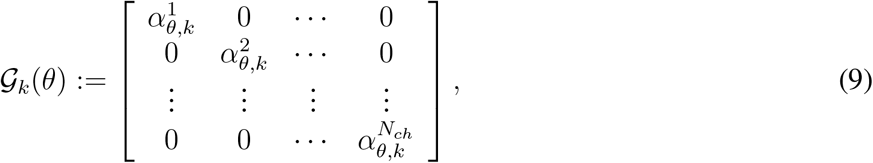

and 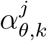 is given by eq. 6 for *j* = 1, …, *N_ch_*, *k* = 1, …, *N_p_* and *θ* ∈ Θ.
2. Let *N_ch_* = *N_s_*. For *θ* ∈ Θ and *k* = 1, …, *N_p_*, let *ν_θ,k_* = *θ_k_*. Then the Fisher information matrix for the output image cube *θ* is given by

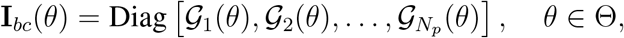

where *𝒢 _k_*(*θ*) is given by eq. 9 for *θ* ∈ Θ and *k* = 1, …, *N_p_*.

**Proof:**

1. By definition of *ν_θ_k*, we have for *θ* ∈ Θ, *j* = 1, …, *N_ch_* and *k* = 1, …, *N_p_*,

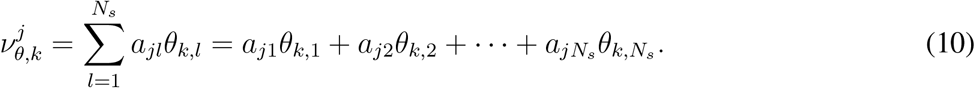 Hence for *k ≠m* and *k, m* = 1, …, *N_p_*, we have

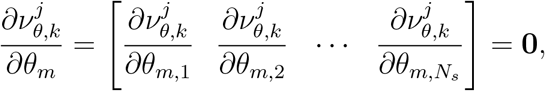

for *θ* ∈ Θ and *j* = 1, …, *N_ch_*, and 0 denotes a 1 × *N_s_* vector with all elements equal to zero. Hence the Fisher information matrix will be given by result 2 of Theorem 2.1. To prove the desired result, it is sufficient if we show that **I**_*k*_(*θ*) = *A^T^ 𝒂 _k_*(*θ*)*A*, where **I**_*k*_(*θ*) is defined by eq. 7. Substituting eq. 10 in eq. 7, we have

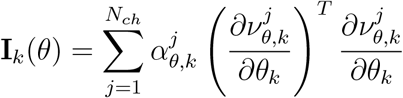

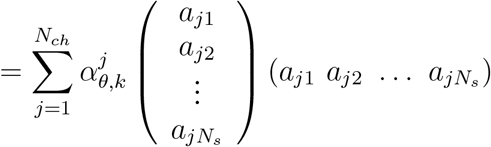

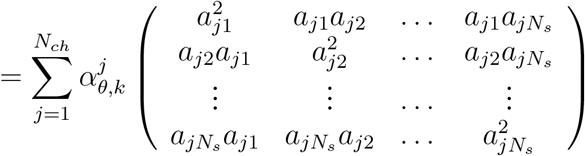

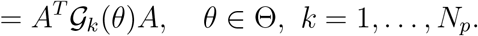
2. Substituting *ν_θ,k_* = *θ_k_* for *θ* ∈ Θ and *k* = 1, …, *N_p_* in eq. 7 and simplifying the result follows.

From result 1 of the above theorem we see that the Fisher information matrix for the linear mixing model depends on the mixing matrix A and the term 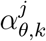 which is given by eq. 6. Note that the analytical expression of 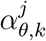 depends on the probability density function of the data model that describes the input image cube. In the next Corollary we derive analytical expression for three of the data models that we discussed in section II-C.

From result 2 of the above Theorem, the Fisher information matrix **I**_*bc*_(*θ*) can be considered as a special case of result 1 with the mixing matrix being equal to a *N_s_ ×N_s_* identity matrix. As we will see, this result will pertain to the best case imaging scenario in that it provides the best possible limit to the accuracy with which photons from a label can be estimated in the output image cube. We note that a hyperspectral imaging system with *A* = 1 may not be practically realizable, for example, if there is significant spectral overlap between the labels used in the sample. Nevertheless, this provides an important benchmark to assess the effect of spectral overlap on spectral unmixing accuracy.

#### Corollary 2.1

For *θ* ∈ Θ, let **I**(*θ*) denote the Fisher information matrix given by result 1 of Theorem 2.2, and for *k* = 1, …, *N_p_* let *ν_θ,k_* be given by eq. 8.

1. **Gaussian data model**. For *θ* ∈ Θ, *j* = 1, …, *N_ch_* and *k* = 1, …, *N_p_*, let 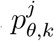 be given by eq. 2. Then for *θ* ∈ Θ

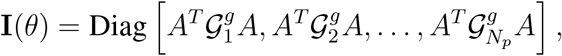

where for *k* = 1, …, *N_p_* and *θ* ∈ Θ

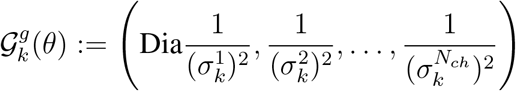

and 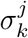 denotes the standard deviation of Gaussian noise for *j* = 1, …, *N_ch_* and *k* = 1, …, *N_p_*.
2. **Poisson data model**. For *θ* ∈ Θ, *j* = 1, …, *N_ch_* and *k* = 1, …, *N_p_*, let 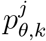 be given by eq. 3. Then for *θ* ∈ Θ

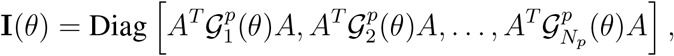

where for *k* = 1, …, *N_p_* and *θ* ∈ Θ

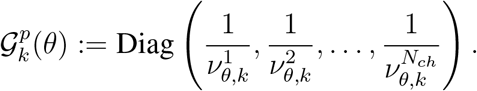
*3.* **Poisson + Gaussian data model**. For *θ* ∈ Θ, *j* = 1, …, *N_ch_* and *k* = 1, …, *N_p_*, let 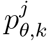 be given by eq. 4. Then

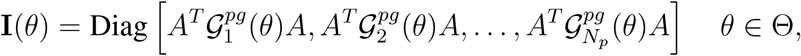

where for *θ* ∈ Θ, *j* = 1, …, *N_ch_* and *k* = 1, …, *N_p_*

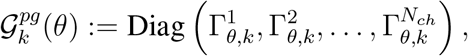

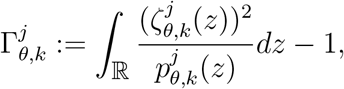

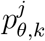 is given by eq. 4 and

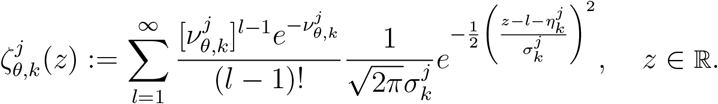

**Proof:**

1. Substituting for 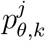 (eq. 2) in eq. 6, we have

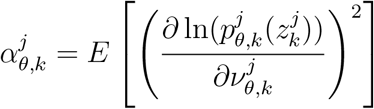

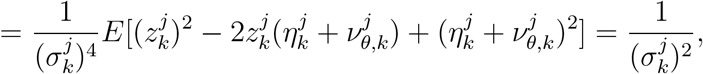

where *θ* ∈ Θ, *j* = 1, …, *N_ch_* and *k* = 1, …, *N_p_*. In deriving the above result we have made use of the fact that the random variables 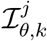 are mutually independent of each other, 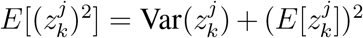 and 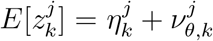 for *θ* ∈ Θ, *k* = 1, …, *N_p_* and *j* = 1, …, *N_ch_*. Substituting the above result in eq. 9 the result follows.
2. For an independent Poisson random variable with mean 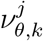, where *θ* ∈ Θ, *j* = 1, …, *N_ch_* and *k* = 1, …, *N_p_*, we have

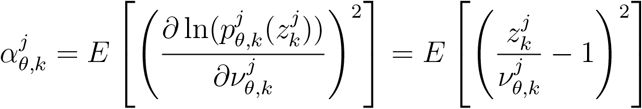

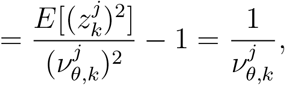

where *θ* ∈ Θ, *j* = 1, …, *N_ch_* and *k* = 1, …, *N_p_*. Substituting the above expression in eq. 9 the result follows.
3. Substituting eq. 4 in eq. 6 and simplifying (see [22] for details), we have

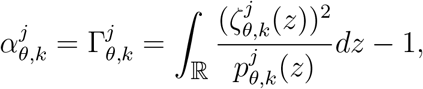

for *θ* ∈ Θ, *j* = 1, …, *N_ch_* and *k* = 1, …, *N_p_*. Substituting the above expression in eq. 9 the result follows.

From the above Corollary we see that the Fisher information matrix for the Gaussian data model is independent of the term 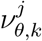 and depends only on the mixing matrix A and the variance 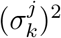 of the Gaussian noise for *θ* ∈ Θ, *j* = 1, …, *N_ch_* and *k* = 1, …, *N_p_*. This is in contrast to the Poisson and the Poisson + Gaussian data models where the Fisher information matrix depends on 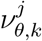 which describes the expected number of detected photons at the *k^th^* pixel in the *j^th^* channel for *θ* ∈ Θ, *j* = 1, …, *N_ch_* and *k* = 1, …, *N_p_*. An implication of this result is that if the underlying data model is indeed Gaussian then the Fisher information matrix of the output image cube is solely governed by the optical configuration of the imaging system and is independent of the photon budget. We note that the above result for the Gaussian data model is consistent with a previously published result ([19]).

From result 2 of Corollary 2.1, we see that the Fisher information matrix for the Poisson data model shows inverse dependence on the expected photon count. Consequently, the inverse Fisher information matrix will show linear dependence on the expected photon count. According to the Cramer-Rao inequality, for an unbiased estimator *θ̂* of an n-dimensional vector parameter *θ*, we have *Var*(*θ̂ i*)≥ [**I**^−1^(*θ*)]_*ii*_, where [**I**^−1^(*θ*)]_*ii*_ denotes the Cramer-Rao lower bound of *θ̂_i_* for *i* = 1, …, *n*. An implication of this result is that the Cramer-Rao lower bound of the photon count estimate for the labels will increase with increasing value of the expected photon count. Since the (square root of) Cramer-Rao lower bound provides a lower bound to the variance (or standard deviation) of the photon count estimate, we will use the ratio of the square root of the Cramer-Rao lower bound of photon count to the expected photon count as a performance measure of the spectral unmixing problem (See Definition 4.2 in Results section).

### F. Channel addition

An important question that arises in the design of hyperspectral imaging systems concerns the number of spectral channels that is required to capture the signal from the label of interest in order achieve optimal spectral unmixing. Specifically, given an input image cube with *N_ch_* spectral channels the question arises as to whether adding an extra spectral channel will provide any benefit for spectral unmixing. In the next Corollary we show that for the linear mixing model with *N_s_* labels, the Fisher information matrix for the output image cube pertaining to a *N_ch_* + 1 channel hyperspectral imaging system is greater than that of the Fisher information matrix pertaining to a *N_ch_* channel hyperspectral imaging system.

#### Corollary 2.2

For *θ* ∈ Θ, let **I**(*θ* | *N_ch_*) denote the Fisher information matrix given by result 1 of Theorem 2.2 pertaining to an input image cube with *N_ch_* spectral channels. Then for *θ* ∈ Θ, **I**(*θ* | *N_ch_* + 1) *≥* **I**(*θ | N_ch_*).

**Proof:**

By definition of **I**(*θ | N_ch_*), it is sufficient to show that **I**_*k*_(*θ | N_ch_* + 1) *≥* **I**_*k*_(*θ | N_ch_*) for *θ* ∈ Θ and *k* = 1, …, *N_p_*, where **I**_*k*_(*θ | N_ch_*) = *A^T^ 𝒂 _k_*(*θ*)*A* for *θ* ∈ Θ, *A* is a *N_ch_* × *N_s_* matrix and *𝒂 _k_*(*θ*) is a *N_ch_* × *N_ch_* diagonal matrix, for *k* = 1, …, *N_p_* that is given by eq. 9. In the case of a *N_ch_* + 1 channel hyperspectral imaging system, we have for *θ* ∈ Θ and *k* = 1, …, *N_p_*

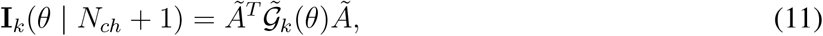

where *Ã* is (*N_ch_* + 1) × *N_s_* matrix and *𝒢̃ k*(*θ*) is (*N_ch_* + 1) *×* (*N_ch_* + 1) diagonal matrix, for *k* = 1, …, *N_p_*. Since the number of labels is the same, we rearrange the mixing matrix *Ã* such that the addition of an extra spectral channel results in the addition of a row at the bottom of this matrix. Hence *Ã* can be written as 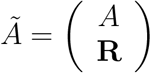 where R is a 1 × *N_s_* vector that pertains to the (*N_ch_* + 1)th channel. Similarly, the matrix *𝒢̃ _k_*(θ) can be written as

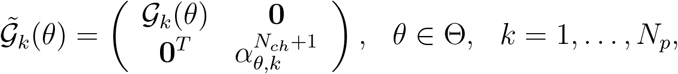

where 0 denotes a *N_ch_ ×* 1 zero vector and 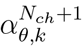 is a scalar term given by eq. 6 and pertains to the (*N_ch_* + 1)th diagonal entry of *𝒢̃_k_* (*θ*). Substituting the above expressions of *𝒢̃_k_* (*θ*) and *A ̃* in eq. 11, we have

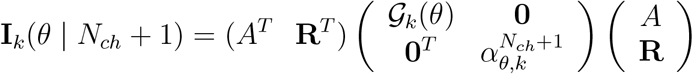

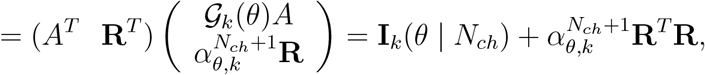

where *θ* ∈ Θ and *k* = 1, …, *N_p_*. By definition, 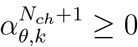 for *θ,* ∈ Θ and *k* = 1, …, *N_p_* and **R**^*T*^ **R** *≥* 0. From this the result follows.

In deriving the above result, we made no specific assumptions about the underlying data model that describes the input image cube. Therefore the above result is applicable to a wide variety of imaging configurations for the linear mixing model. An immediate implication of the above result is that the limit of the accuracy of photon count for a given label improves by adding extra spectral channels to detect the signal from that label.

### G. Channel splitting and photon partitioning

In the previous section, we saw how adding an extra channel to a hyperspectral imaging system can improve the Fisher information matrix. In many practical situations, channel addition may not be feasible since the overall spectral bandwidth to collect the signal from a given label cannot be increased. In such cases, a question then arises as to whether spectral subsampling within the passband would be beneficial. Specifically, if we have a hyperspectral imaging system with *N_ch_* spectral channels and if one of the spectral channels are split into two channels of smaller spectral bandwidth, then under what conditions is channel splitting beneficial? In the next corollary we show that channel splitting is beneficial for the Poisson data model.

#### Corollary 2.3

For *θ* ∈ Θ, let **I**(*θ | N_ch_*) denote that Fisher information matrix given by result 2 of Theorem 2.1 where *N_ch_* denotes the number of spectral channels. Assume that for *j* = 1, …, *N_ch_* there exists a partition function 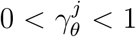 with 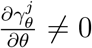 for *θ* ∈ Θ and *j* = 1, …,*N _ch_* such that the addition of an extra channel splits the expected photon counts into two fractions given by 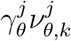 and 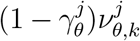, where 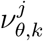 denotes the expected photon count of the *k^th^* pixel in the *j^th^* spectral image, for *θ* ∈ Θ, *j* = 1, …, *N_ch_* and *k* = 1, …, *N_p_*. If we consider a Poisson data model, then

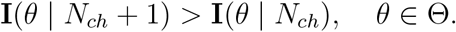

**Proof:**

By definition of **I**(*θ | N_ch_*), it is sufficient to show that **I**_*k*_(*θ | N_ch_* + 1) *≥* **I**_*k*_(*θ | N_ch_*) for *θ* ∈ Θ and *k* = 1, …, *N_p_*, where **I**_*k*_(*θ | N_ch_*) is given by eq. 7. Without loss of generality, we set *N_ch_* = 1. For a Poisson data model, it can be shown that 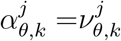 for *θ* ∈ Θ, *j* = 1, …, *N_ch_* and *k* = 1, …, *N_p_* where 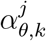 is defined in eq, 6. Substituting this in eq. 7, we get

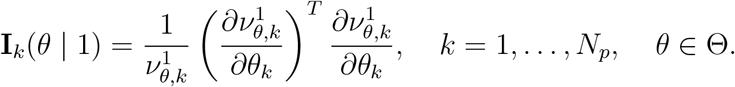

If a second spectral channel is introduced (*N_ch_* = 2) and a fraction of the photons detected in the first channel is now captured in the second channel, then we have for *θ* ∈ Θ and *k* = 1, …, *N_p_*,

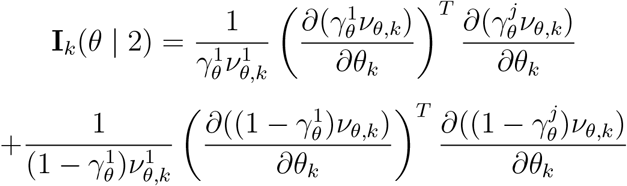

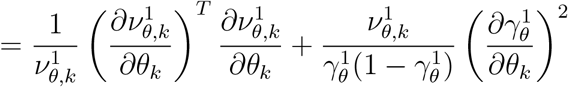

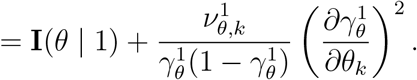

From this the result follows.

Note that unlike Corollary 2.2 which used the analytical expression for the Fisher information matrix given in Theorem 2.2, the above corollary used a more general expression for the Fisher information matrix given by result 2 of Theorem 2.1. The reason for this is due to the dependence of *θ* by the partition function 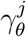 for *j* = 1, …, *N_ch_* and *θ* ∈ Θ. As we will show in Section IV-E, the above result is only true for the Poisson data model. In fact for the Poisson + Gaussian data model we show that the limit of the accuracy of the photon count exhibit complex behavior with channel splitting and eventually deteriorates when increasing the number of partitioned channels. We note that the result given in Corollary 2.3 was also reported in [20] for the Poisson data model.

## III. Methods

### A. Computing the Fisher information matrix and the mixing matrix A

The Fisher information matrix was calculated using the FandPLimitTool ([27]) which is a MATLAB based software package that contains an extensive suite of tools to calculate the Fisher information matrix for a wide variety of parameter estimation problems in fluorescence microscopy. For the linear spectral unmixing problem, the Fisher information matrix depends on the mixing matrix A which needs to be computed for for illustrating the results. We note that in a practical situation, *A* will be experimentally determined by imaging single color control samples under identical imaging conditions to that of the sample that contains *N_s_* different labels. Here we present a simplified approach to theoretically calculate A by using relevant reference data such as the the excitation and emission spectra of fluorescent labels, the spectral emissivity of the fluorescent light source and passbands of the excitation and emission filters. The elements of the mixing matrix A can be written as

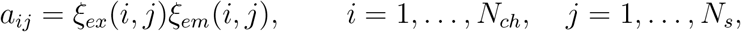

where for *i* = 1, …, *N_ch_* and *j* = 1, …, *N_s_*

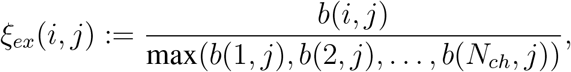

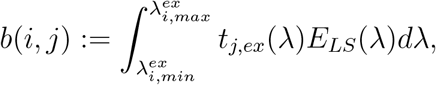

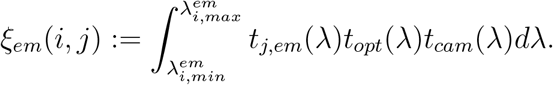

In the above equation 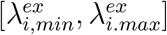 and 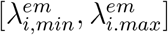 denote the excitation and emission passbands corresponding to the *i^th^* channel, *t_j,ex_* and *t_j,em_* denote the fluorescence excitation and emission spectra of the *j^th^* dye, *j* = 1, …, *N_s_*, *E_LS_* denotes the spectral emissivity of the fluorescent light source, *t_opt_* denotes the spectral transmitivity of the optical components (e.g., objective and tube lens) in the emission light path, and *t_cam_* denotes the spectral sensitivity of the detector. For simplicity, we set *t_opt_*(*λ*) = 1 and *t_cam_*(*λ*) = 1, and also assume that the transmission efficiency of the excitation and emission filters is equal to 1 in the passbands and equal to zero outside the passbands. The excitation and emission spectra of the fluorescent labels and the spectral emissivity of the fluorescent light source were obtained from the Pubspectra database [28] and were modified such that their sum equals 1. The above integrals were numerically evaluated using the Trapezoidal rule.

### B. Hyperspectral data simulation

All simulations and estimations were carried out in the MATLAB programming language. To simulate hyperspectral data, we use the above approach to compute the mixing matrix A and use eq. 8 to compute the expected photon counts of the fluorescent labels *ν_θ,k_* at the k^*th*^ pixel in the input image cube. For the Poisson + Gaussian data model, we first create a Poisson realization of *ν_θ,k_* and then add Gaussian noise with mean 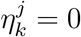 e^−^/pixel and standard deviation 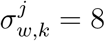 e^−^/pixel to the Poisson data.

## IV. Results

In this section, we illustrate the results obtained in the previous section concerning the linear mixing model. We first define a few entities that we will use throughout this section.

### Definition 4.1

For *θ* ∈ Θ, *i* = 1, …, *N_s_* and *k* = 1, …, *N_p_*, the **Linear Unmixing Performance (LUP) bound** for the i^*th*^ label at the *k^th^* pixel in the output image cube is defined as 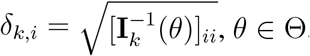, and the **LUP bound for the best case scenario** is defined as 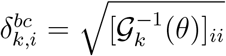, where **I**_*k*_(*θ*) = *A^T^ 𝒢 _k_*(*θ*)*A*, *A* denotes the mixing matrix and *𝒢 _k_*(*θ*) is given by 9, for *θ* ∈ Θ and *k* = 1, …, *N_p_*.

### Definition 4.2

For *θ* ∈ Θ, *i* = 1, …, *N_s_* and *k* = 1, …, *N_p_*, the **normalized LUP (nLUP) bound** for the i^*th*^ label at the *k^th^* pixel in the output image cube is defined as 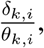 and the **nLUP bound for the best case scenario** is defined as 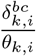, where *δ_k,i_* and 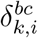 are defined in Definition 4.1.

### A. Spectral resolution for a pair of fluorophores

An important question that arises in hyperspectral imaging applications concerns with how accurately two spectrally overlapping fluorescent labels can be discerned from the acquired data. Here we address this problem by considering a pair of fluorescent labels whose emission maxima are separated by a distance *d_s_* (see Fig. 2A). We assume an imaging configuration where the sample is illuminated (sequentially or simultaneously) to excite the labels such that the excitation passband for a given label overlaps with its corresponding excitation maxima. The fluorescence signal is then detected (sequentially or simultaneously) in two distinct spectral channels, where the emission passband of each spectral channel covers the corresponding emission maxima of a particular label. We then ask the question how decreasing the spectral distance *d_s_* affects the spectral resolution of the two fluorophores. For the current discussion, we consider the fluorescent labels with traditional Stokes shift fluorescence signal ([29]) where the peak of the emission spectra for the label occurs at a longer wavelength (lower energy) than the peak of its corresponding excitation spectra.

**Fig. 2.**
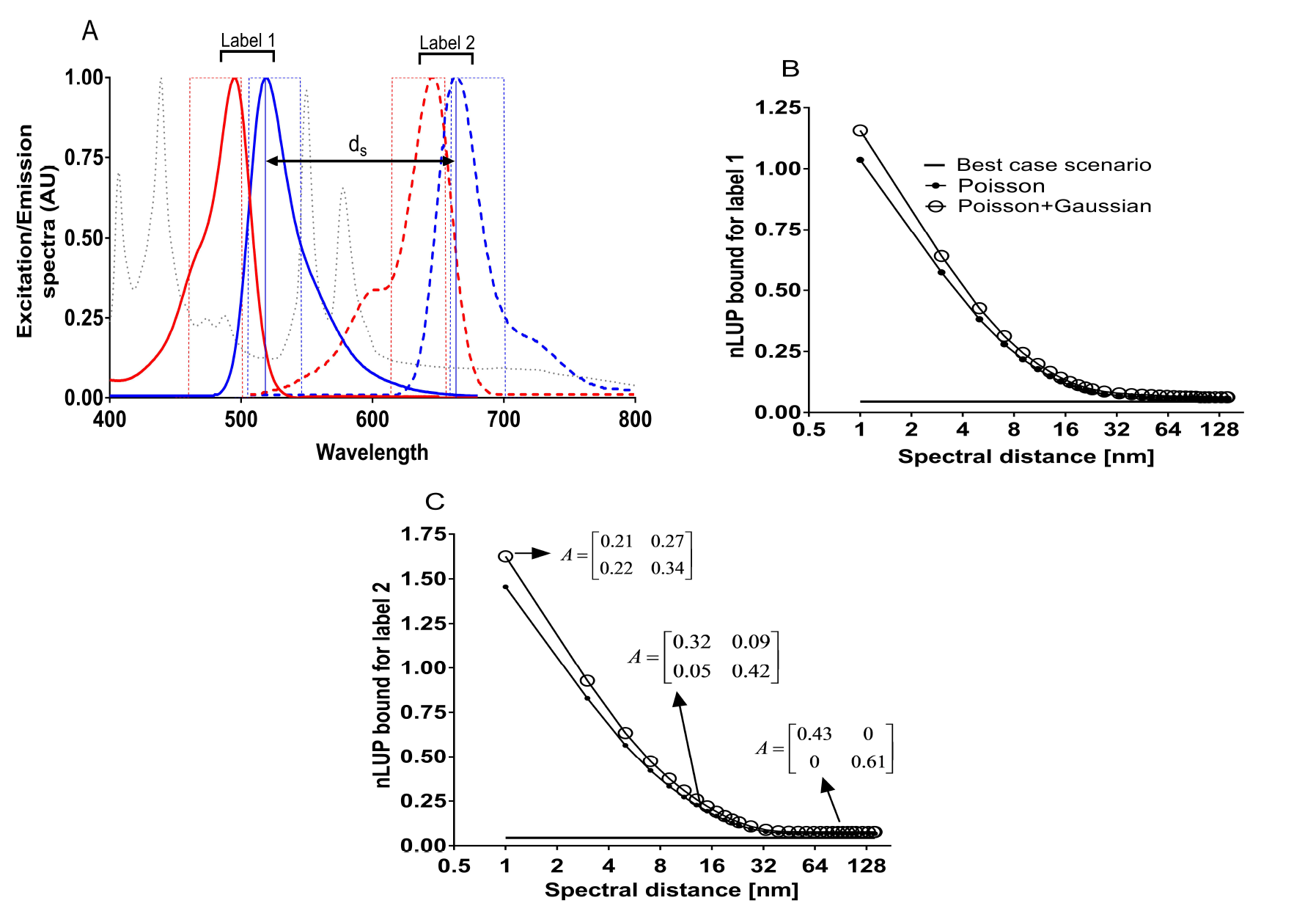
The effect of changing the spectral distance on spectral resolution. Panel A shows the normalized excitation (red lines) and emission spectra (blue lines) of two fluorescent labels that are spectrally separated by a distance *d_s_*. The panel also shows the spectral emissivity of a metal hallide broadband light source that is used to excite the two labels. Here, for both labels we consider the traditional Stoke’s shift for their fluorescence emission spectra. Panels B and C show the nLUP bound for labels 1 and 2 at a pixel, respectively, for the Poisson and the Poisson + Gaussian data models and also for the best case scenario pertaining to the Poisson data model. Panel C also shows the mixing matrix pertaining to different values of *d_s_*. In panels B and C, the expected photon count for both labels is set to be 500 photons per pixel and for the Poisson + Gaussian data model we consider the mean and standard deviation of the Gaussian noise to be 0 e^*−*^/pixel and 8 e^*−*^/pixel, respectively.

Figs. 2B and 2C show the nLUP bounds for labels 1 and 2, respectively, for different data models as a function of the spectral distance *d_s_*. The figures also show the nLUP bounds for labels 1 and 2 for the best case scenario, which pertains to the ideal imaging configuration wherein the signal collected from a given label is not corrupted by photons from other labels present in the sample. Here, to simulate the decrease in *d_s_*, we shift the excitation and emission spectra of label 2 and also the corresponding passbands of the excitation and emission filters towards label 1 while leaving the excitation and emission spectra for label 1 unchanged. By definition, the nLUP bound is a ratio of the LUP bound to the expected photon count for a given label (Definition 4.2). A large numerical value of the nLUP bound predicts a relatively high level of error in estimating the photon count at a given pixel while a small numerical value predicts a relatively low level of error in estimating the photon count at a given pixel.

From the figures we see that when the labels are spectrally well separated the nLUP bound is numerically close to the nLUP bound for the best case scenario. This is expected since there is no spectral crosstalk between the excitation or emission spectra of the two labels. Consistent with this, we see that the mixing matrix A is diagonal (see Fig. 2C). As the spectral distance decreases, the excitation/emission spectra of the labels start to overlap and the mixing matrix is no longer diagonal. Consequently, the numerical value of the nLUP bound becomes bigger and starts to deviate from the nLUP bound for the best case scenario. When the spectral distance further decreases, the nLUP bound continues to deteriorate and we see that the numerical values of the off-diagonal terms of the A matrix, which accounts for the spectral cross talk between the labels, is comparable to the diagonal terms. Note that the nLUP bound for the Poisson + Gaussian data model is consistently higher than that for the Poisson data model, since in the former model the presence of additive Gaussian noise corrupts the detected photon counts at each pixel in the input image cube which leads to higher uncertainty in estimating the photon counts at a given pixel.

We note that the difference between the nLUP bound and the nLUP bound for the best case scenario is referred to as the multiplexing disadvantage ([15]) which accounts for the deterioration in the limit of the accuracy of photon count from the best case scenario when we consider the effect of detecting unwanted photons from other labels present in the sample. As we will see in rest of this section, the nLUP bound for the best case scenario will provide an important benchmark for studying the performance limits of hyperspectral imaging systems. In our present example the shape of the emission spectra for labels 1 and 2 are dissimilar, and therefore even for a very small spectral distance of 1 nm, the nLUP bound remains finite for both labels.

### B. Improving the spectral resolution by using anti-Stokes shift fluorescence

In the previous section we saw that with traditional fluorescent labels which exhibit Stokes shift fluorescence, there is a limit to how spectrally close the two labels can be and still be accurately spectrally unmixed. In this section, we show that by replacing one of the labels with a fluorophore that exhibits anti-Stokes shift fluorescence we can overcome the spectral resolution limit. Fig. 3 shows the excitation and emission spectra for such a pair of labels, where label 1 is a traditional fluorescent label as in Fig. 2 and label 2 exhibits anti-Stokes shift fluorescence ([29]) wherein the peak of its emission spectra occurs at a shorter wavelength (higher energy) relative to the peak of its excitation spectra. It should be pointed out that at present there is significant interest in developing such probes especially for biological applications ([30], [31], [32]). For example, upconverting nanoparticles are one such a class of fluorescent labels that typically exhibit strong absorption in the near infrared wavelength range (700 - 1000 nm) and emit fluorescence signal in the visible range (400 - 650 nm) ([33], [34]).

**Fig. 3.**
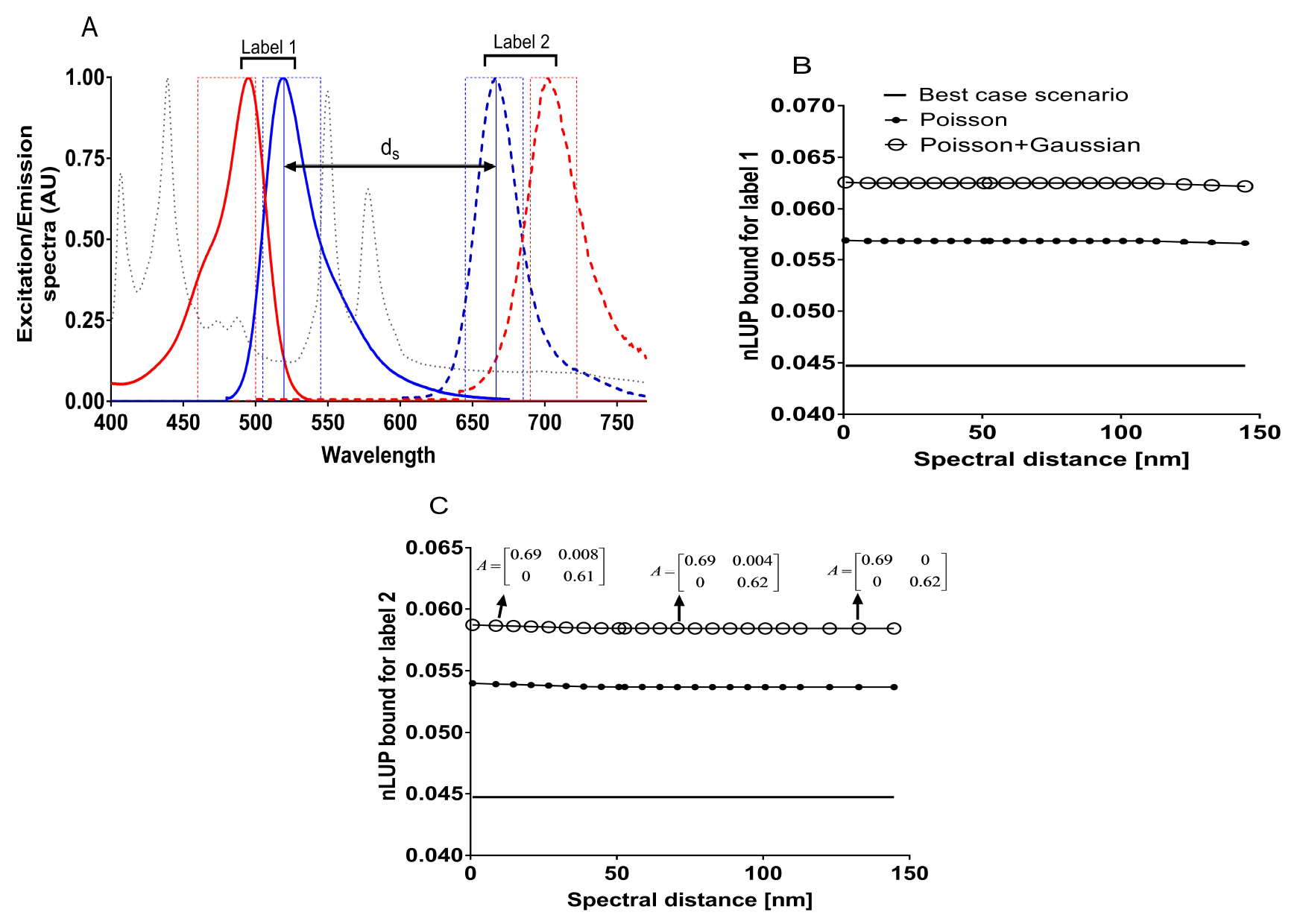
Improving the spectral resolution by using anti-Stokes shift fluorescences. Panel A shows the normalized excitation (red lines) and emission spectra (blue lines) of two fluorescent labels that are spectrally separated by a distance *d_s_*. The panel also shows the spectral emissitivity of a metal hallide broadband light source that is used to excite the two labels. Here, label 1 is the same as that shown in Fig. 2 while label 2 is a fluorophore with anti-Stokes shift fluorescence emission, where the peak of its emission spectra is at a shorter wavelength (higher energy) than the peak of its excitation spectra. Panels B and C show the nLUP bounds for labels 1 and 2, respectively, at a pixel for the Poisson and the Poisson + Gaussian data models and also for the best case scenario pertaining to the Poisson data model. Panel C also shows the mixing matrix pertaining to different values of *ds*. The numerical values used to generate the above plots are identical to those used in Fig. 2.

In fig. 3 we have simulated the excitation and emission spectra for a hypothetical label that exhibits anti-Stokes shift fluorescence. We consider an imaging configuration where the labels are sequentially illuminated over two excitation passbands that cover their corresponding excitation maximas, and the fluorescence signal is sequentially collected over emission passbands that cover their emission maximas. To simulate the decrease in spectral distance, we shift the excitation and emission spectra of label 2 and their corresponding excitation and emission passbands towards label 1, and keep the excitation and emission spectra of label 1 fixed.

Figs. 3B and 3C show the nLUP bounds for labels 1 and 2, respectively, pertaining to the Poisson and Poisson+Gaussian data models. The figures also show the nLUP bound for the best case scenario pertaining to the Poisson data model. From the figures we see that the nLUP is almost constant for all values of the spectral distance. Specifically, the mixing matrix A remains diagonal irrespective of the spectral distance of separation between the two labels. This can attributed to anti-Stokes shift fluorescence which significantly eliminates the overlap of the excitation spectra between labels 1 and 2 when the spectral distance decreases. Therefore for above the imaging configuration there is very limited spectral cross talk between the two labels and consequently the nLUP bound remains constant and is consistently close to the nLUP bound for the best case scenario. Note that similar to Fig. 2, the nLUP bound for the Poisson+Gaussian data model is consistently higher than the nLUP bound for the Poisson data model.

### C. Effect of photon count

We next investigate how changing the expected photon count impacts the linear unmixing performance bound. Here, we consider two fluorescent labels Cy3 and Cy3.5 which have significant overlapping excitation and emission spectra (Figs. 4A and 4B). We assume an imaging configuration in which the fluorophores are sequentially excited at distinct excitation passbands and the corresponding fluorescence signal is then sequentially collected at the indicated emission passbands.

**Fig. 4.**
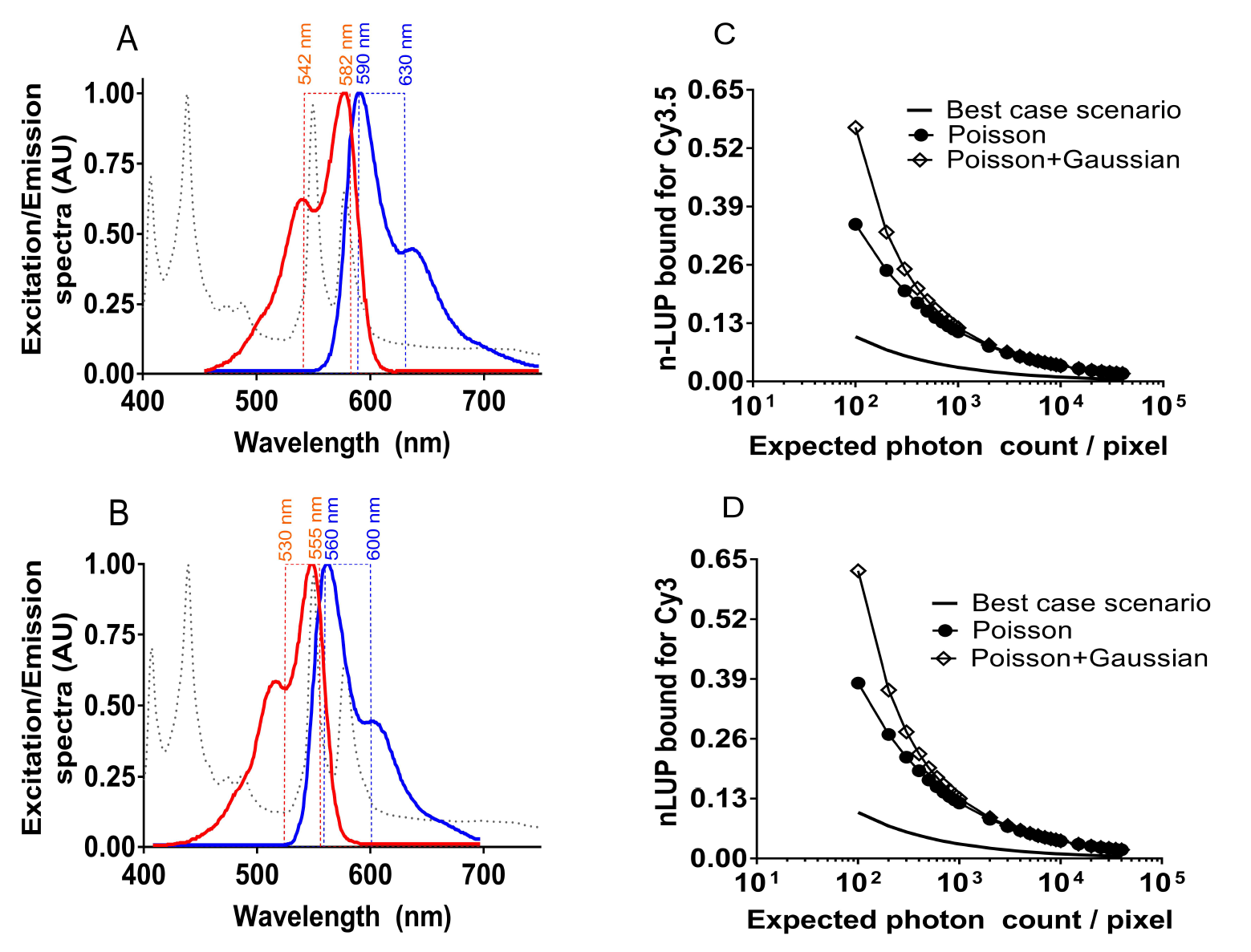
Effect of photon count on the nLUP bound. Panels A and B show the normalized excitation (red lines) and emission (blue lines) spectra for Cy3.5 and Cy3 fluorescent labels, respectively, along with the spectral emissivity of a metal hallide light source (dotted line). The panels also show the corresponding excitation and emission passbands for each fluorescent label. Panels C and D show the behavior of the nLUP bound for Cy3.5 and Cy3, respectively, at a single pixel as a function of the expected photon count for the Poisson and the Poisson + Gaussian data models. As reference, the panel also shows the nLUP bound for the best case scenario pertaining to the Poisson data model. Here, we assume the expected photon count to be the same for both fluorophores and for the Poisson + Gaussian data model, the mean and standard deviation of the readout noise is set to be 0 *e^−^*/pixel and 8*e^−^*/pixel, respectively.

Figs. 4C and 4D show the nLUP bound for the Poisson and the Poisson + Gaussian data models along with the nLUP bound for the best case scenario pertaining to the Poisson data model. We see that as the expected photon count /pixel increases, the numerical value of the nLUP bound becomes smaller and approaches the nLUP bound for the best case scenario. Note that the nLUP bound for the Poisson + Gaussian data model is consistently higher than that of the Poisson data model, and the difference starts to diminish as the expected photon count / pixel increases. Taken together, these results imply that increasing the photon/light budget would result in an overall improvement in the performance bound.

### D. Channel addition

We next investigate the effect of adding extra spectral channels to collect the signal from a label on the spectral unmixing accuracy. Here again we consider Cy3 and Cy3.5 labels and calculate the LUP bound that is defined in Definition 4.1. Unlike the results given in Figs. 3 and 4, here we compute the LUP bound since the expected photon count is fixed for the fluorescent labels. We assume an imaging configuration such that the fluorescence signal from each label is sequentially collected in one or more spectral channels (i.e. different emission passbands) by repetitively exciting the sample at the same excitation passband. For example, for the Cy3.5 fluorophore, consider a 6 channel configuration with emission passbands in the range of 590 - 610 nm, 610 - 630 nm, 630 - 650 nm, 650 - 670 nm, 670 - 690 nm and 690 - 710 nm (see Fig. 5A). Here the sample will be sequentially excited 6 times at the excitation passband of 542 - 582 nm and each time the fluorescence signal will be collected in a different emission passband pertaining to Cy3.5. Note that in this configuration the number of detected photons from the fluorescent label will increase with increasing number of channels assuming that there is negligible photobleaching effect which typically diminishes the fluorescence intensity of the label with repeated excitation. We note that such an imaging configuration has been implemented, for example, using specialized emission filters that are placed before the imaging detector in the light path in which the passband of the filter can be changed either electronically ([9]) or optomechanically by changing the relative angle of incidence of light on the emission filter ([7])

**Fig. 5.**
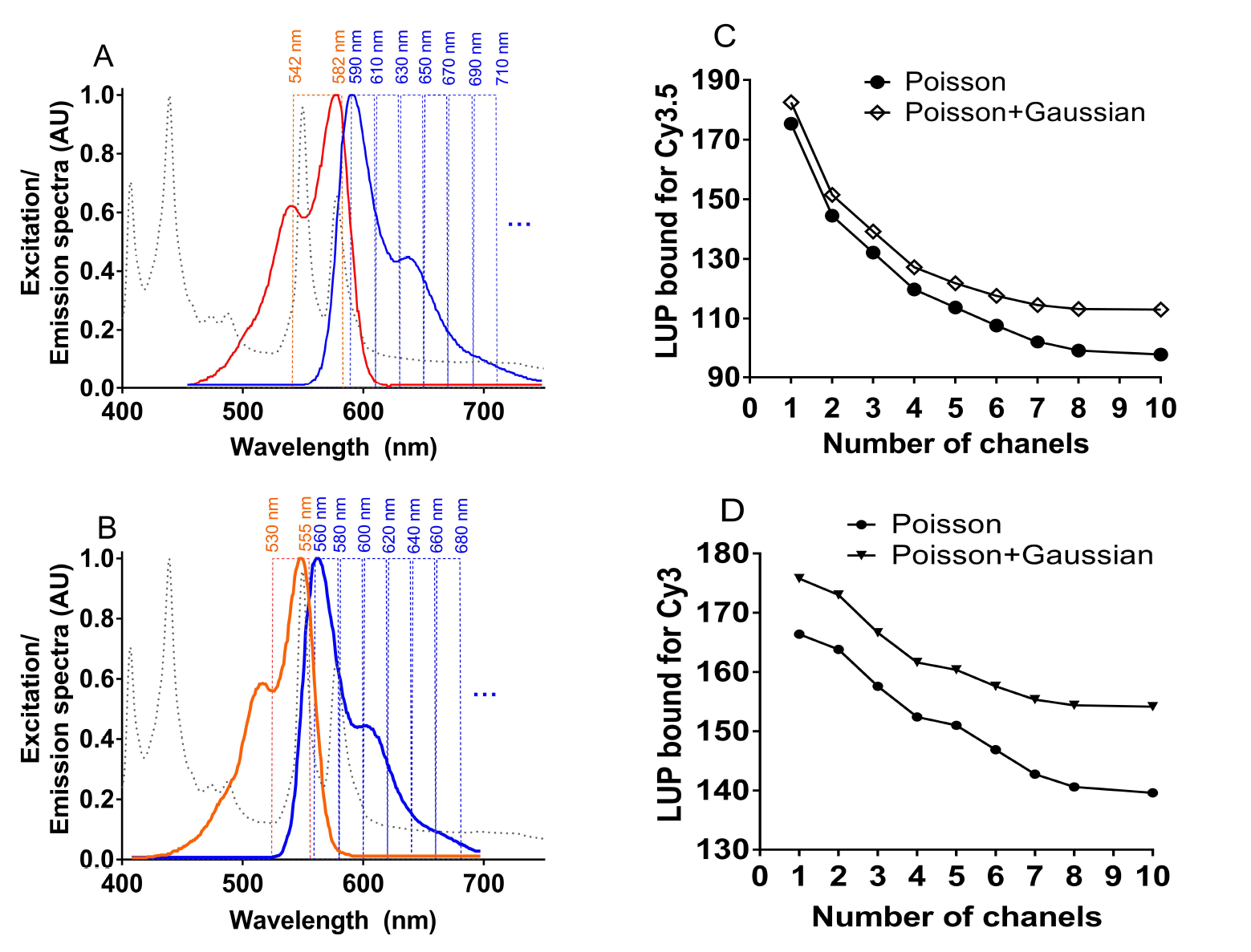
Effect of channel addition on the LUP bound. Panels A and B show the normalized excitation (red lines) and emission (blue lines) spectra for Cy3.5 and Cy3 fluorescent labels, respectively, along with the spectral emissivity of a metal hallide light source (dotted line). The panels also show the excitation passband and the emission passbands pertaining to the different spectral channels. Panel C shows the behavior of the LUP bound for Cy3.5 at a single pixel as a function of the number of channels after channel addition for different data models. Panel D shows the same for Cy3. In Panels C and D, the expected photon count is set to be 3000 for both labels, and the numerical values of the mean and standard deviation of the Gaussian noise component are identical to those used in Fig. 4.

Figs. 5C and 5D show the LUP bound for Cy3.5 and Cy3 labels, respectively, as a function of the number of spectral channels for two different data models. From the figure we see that for both the Poisson and the Poisson + Gaussian data models the LUP bound improves by increasing the number of channels. This is consistent with Corollary 2.2, where we showed that the Fisher information matrix for an output image cube pertaining to an *N_ch_* + 1 channel hyperspectral system is greater than or equal to that of the Fisher information matrix for an *N_ch_* channel hyperspectral imaging system. Note that for both labels, the LUP bound for the Poisson + Gaussian data model is consistently higher than that of the Poisson data model, which is consistent with the behavior that we observed in Fig. 4. We also observe that the addition of spectral channels beyond a certain wavelength range is not beneficial as we see that the LUP bound starts to plateau out.

### E. Channel splitting

In the previous section we investigated how the addition of spectral channels impacts the LUP bound for a given label. In many situations, the addition of spectral channels is not feasible. More specifically the emission passband for a given fluorescent label is typically fixed. In such cases a question arises as to whether splitting the available passband into narrower spectral channels will render any benefit. Here we investigate this question for Cy3 and Cy3.5 labels. For each fluorescent label, we consider a partitioning scheme wherein we begin with a single emission passband and then partition the passband into 2, 4, 8, 13, 20 and 40 smaller bands. For example, for Cy3.5 we start with the emission passband of 590 nm - 630 nm (see Fig. 4), which we split into two narrower channels, i.e. 590 nm - 610 nm and 610 nm - 630 nm, then into 4 narrower channels, i.e. 590 nm - 600 nm, 600 nm - 610 nm, 610 nm - 620 nm and 620 nm - 630 nm, and so on. A similar partitioning scheme is also adopted for Cy3 where we start with the emission passband of 560 - 600 nm.

We consider an imaging configuration where the labels are sequentially excited at a specific excitation passband followed by simultaneous detection of the emitted photons across the different spectral channels for that label. Unlike the imaging configuration that we considered in Section IV-D, here the total number of photons collected from a given label remains the same as we increase the number of partitioned channels. We note that the above imaging configuration can be realized in a spectral confocal microscope or in a hyperspectral line scanning microscope that is equipped with a dispersive optical element such as a prism or a monochromator, which spectrally disperses the incident light along a line and the dispersed signal is then captured on a linear imaging detector.

Figs. 6A and 6B, show the LUP bound for Cy3.5 and Cy3 labels at a given pixel as a function of the number of partitioned channels for different data models. For the Poisson data model we see that channel splitting improves the LUP bound for both labels. Specifically, the LUP bound decreases with increasing number of channels after partitioning. This is consistent with Corollary 2.3, where we showed that for the Poisson data model the Fisher information matrix for a hyperspectral imaging system with *N_ch_* + 1 partitioned channels is greater than that with *N_ch_* partitioned channels.

**Fig. 6.**
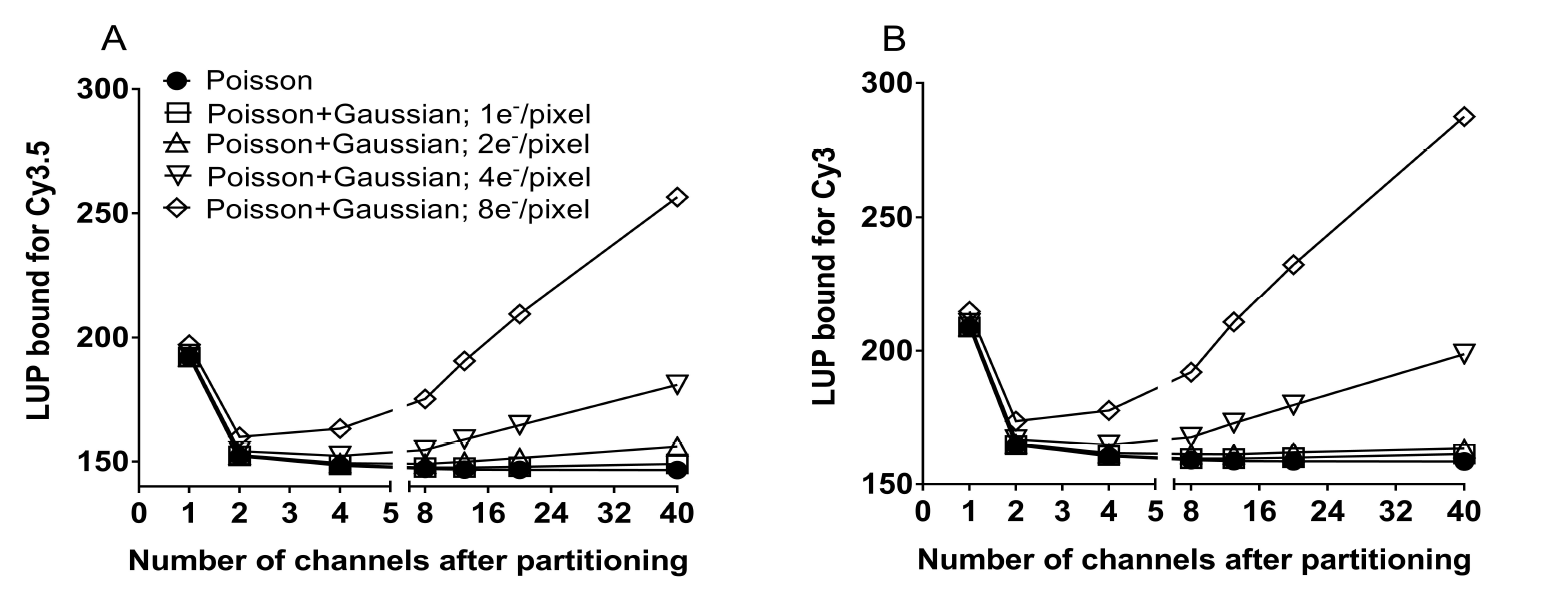
The effect of channel splitting on the LUP bound. Panel A shows the behavior of the LUP bound for Cy3.5 at a single pixel as a function of the number of channels after channel splitting for the Poisson and the Poisson + Gaussian data models. Panel B shows the same for Cy3. In Panels A and B, the expected photon count is set to be 3000 for both labels, and for the Poisson + Gaussian data model we assume the mean of the Gaussian noise to be 0 e^*−*^/pixel and different values of standard deviation as indicated in the legend in panel A.

**Figure 7.**
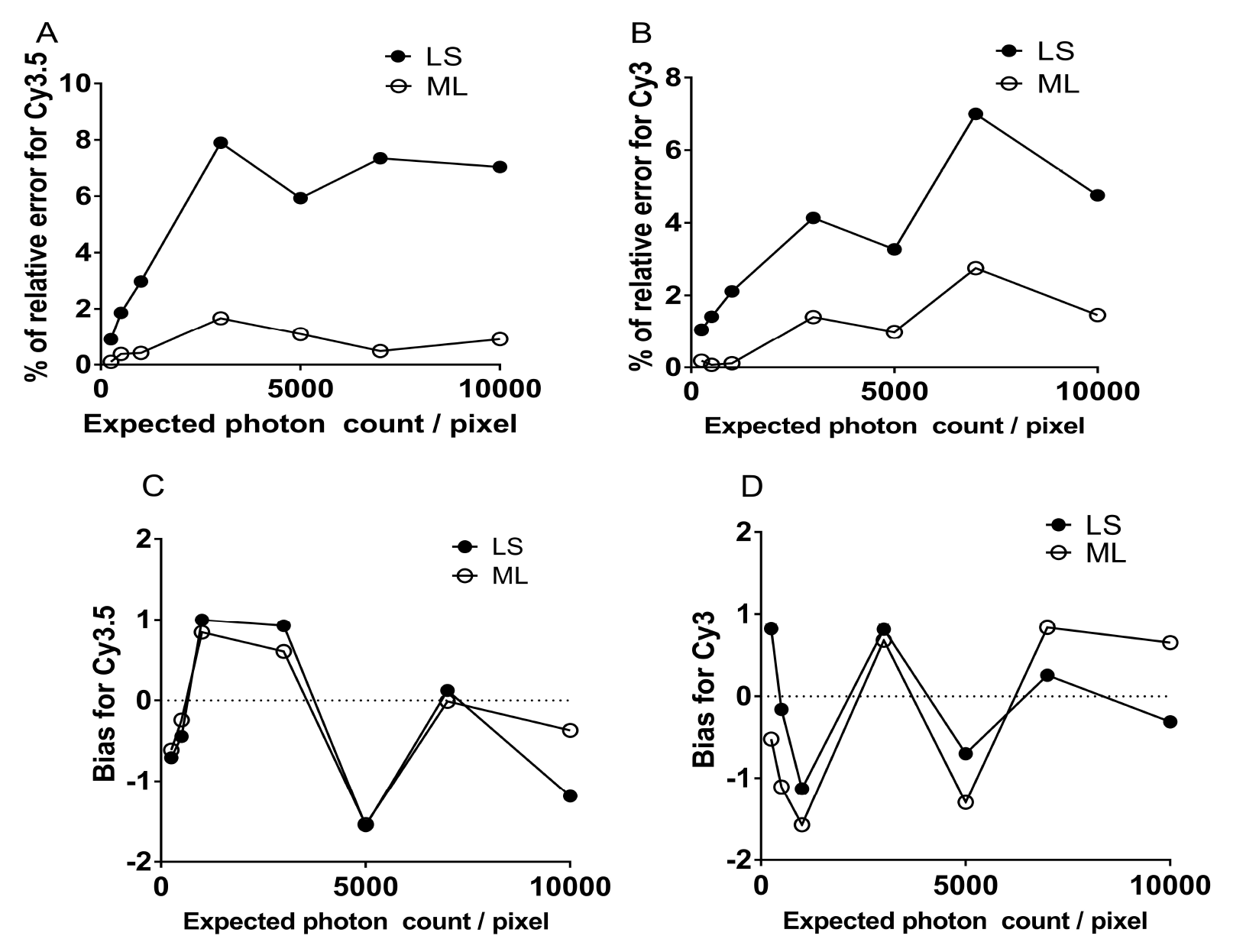
Performance of spectral unmixing algorithms on hyperspectral data. The figure shows the behavior of the LS and ML estimators on simulated hyperspectral data. Panels A and B show the % of relative error of the estimators for Cy3.5 and Cy3 labels, respectively, for different expected photon counts. Here we assume that the expected photon count is the same for both fluorescence labels. Panels C and D show the bias of the estimators for Cy3.5 amd Cy3, respectively, for different expected photon counts.

Figs. 6A and 6B also show the LUP bound for the Poisson + Gaussian data model for different readout noise levels. Here we assume that the readout noise remains the same as the number of partitioned channels increases. Unlike the Poisson data model, we see that the LUP bound first improves but then progressively deteriorates with increasing number of partitioned channels. Specifically, when the number of channels initially increases, this provides additional spectral information which in turn results in an improvement of the LUP bound. However, as the number of partitioned channels increases, the passband for each spectral channel becomes narrower and in turn the expected photon count per spectral channel decreases while the readout noise remains the same.

Note that the extent of this deterioration is proportional to the standard deviation of the readout noise. For example for Cy3.5 label, if the standard deviation of the readout noise is 8 e^*−*^ /pixel, then the LUP bound varies from 160 photons to 210 photons when the number of channels after partitioning increases from 2 to 20. On the other hand, if the standard deviation of the readout noise is 2 e^*−*^/pixel, then the LUP bound varies from 149 photons to 152 photons when the number of channels after partitioning increases from 2 to 20. We note that a similar result has been reported for the localization accuracy problem in fluorescence microscopy, where the limit of the accuracy of the X/Y location coordinate also shows an analogous behavior as a function of pixel size of the imaging detector for the Poisson and the Poisson + Gaussian data models ([35]).

## V. Assessment of Spectral Unmixing Algorithms

In the previous sections we investigated the behavior of the linear unmixing performance bound for different experimental configurations and imaging conditions. A fundamental question remains whether there exists an unbiased estimator than will attain the performance bound. To address this question, we consider two widely used algorithms for spectral unmixing namely the least squares (LS) estimator and the maximum likelihood (ML) estimator, and evaluate their performance on simulated hyperspectral imaging data. Recall from Theorem 2.2 that the block diagonality of the Fisher information matrix implies that the accuracy with which the photon counts are estimated at the k^*th*^ pixel in the output image cube is independent of the accuracy of photon count estimates at the m^*th*^ pixel when *k ≠ m*. Hence for the current discussion it is sufficient to evaluate the performance of the spectral unmixing algorithms for hyperspectral data simulated at a single pixel. For *k* = 1, …, *N_p_*, let 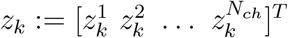, denote the detected photon counts at the k^*th*^ pixel block in the input image cube. The LS estimator of *θ_k_* can be written as

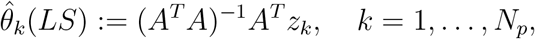

where *A* denotes the mixing matrix. The ML estimator can be written as

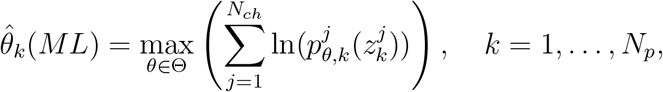

where 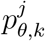 denotes the probability density function of the photon counts detected at the *k^th^* pixel in the *j^th^* spectral channel, for *j* = 1, …, *N_ch_* and *k* = 1, …, *N_p_*.

From the definition it is straightforward to see that for the Poisson data model (eq. 3) the LS estimator is unbiased, since

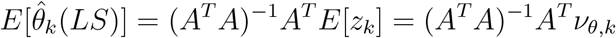

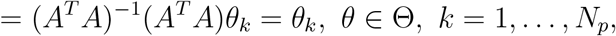

where we have used eq. 8. However, if the hyperspectral data follows either the Gaussian data model (eq. 2) or the Poisson + Gaussian data model (eq. 4), then we have

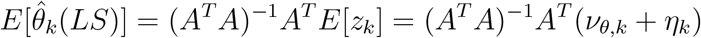

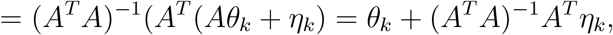

where 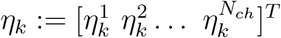 and *k* = 1, …, *N_p_*. From the above equation we see that the LS estimator will be unbiased only when the mean of the Gaussian noise component 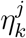 is equal to zero for *j* = 1, …, *N_ch_* and *k* = 1, …, *N_p_*.

We consider the hyperspectral imaging configuration illustrated in Fig. 5 where the fluorescent labels Cy3 and Cy3.5 are sequentially excited and their fluorescence signal is detected across 6 distinct spectral channels. We assume that the expected photon count for each label is the same and simulate the hyper-spectral data as described in Section III-B. For each value of expected photon count, we create 10,000 realizations of the data and estimate the photon counts associated with each fluorescent label by using the different spectral unmixing algorithms. To assess the performance of the unmixing algorithms, we calculate the bias and the relative error which are given by

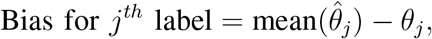

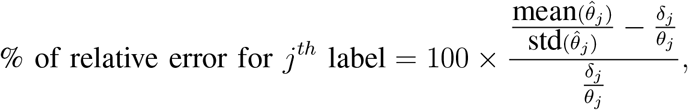

where *j* = 1, 2, *δ_j_/θ_j_* denotes the nLUP bound for *θ_j_* (see Definition 4.2).

Fig. V shows the performance of the LS and ML estimators for the hyperspectral data generated for different values of the expected photon counts. From the figure, we see that the performance of LS and ML estimators are relative close to the nLUP bound in that the % of relative error for both labels are within 8% of their corresponding nLUP bounds. Note that the performance of the ML estimator is consistently better than that of the LS estimator for a range of expected photon count values, since the % of relative error for the ML estimator is lower than that for the LS estimator. This is expected since we used the Poisson + Gaussian data model to simulate the hyperspectral data and used the appropriate probability density function to estimate the photon counts in the ML estimator. The figure also shows the bias for the LS and ML estimators. By definition the LS estimator is unbiased and consistent with this we see that the bias estimates of the LS estimator for both labels are equally distributed between positive and negative values. In particular, the magnitude of the bias estimate is within *±* 1 photon from the true value of the photon count. An almost identical behavior is observed for the ML estimator which suggests that the ML estimator is also unbiased for the imaging conditions tested here.

## VI. Conclusion

A general stochastic model for hyperspectral imaging data was presented and analytical expressions for the Fisher information matrix was derived. The model is based on relatively broad assumptions about the underlying probability density function that describes the hyperspectral data and allows for a wide variety of imaging conditions. As an application, we considered the linear mixing model and showed that the Fisher information matrix becomes block diagonal. Using the Cramer-Rao inequality, we introduced a linear unmixing performance bound and showed how this can be used to predict the spectral resolution limit for two spectrally overlapping fluorescent labels. We also showed how the spectral resolution limit can be surpassed in a standard microscope configuration by using the phenomenon of anti-Stokes shift fluorescence. Further, we investigated the effects of channel addition and channel splitting on the the behavior of the performance bound and illustrated them through concrete examples. Finally, we evaluated the performance of the least squares and maximum likelihood estimators for spectral unmixing, and studied their bias and variance behaviors at different photon/light budgets. In conclusion we note that the results and analysis presented here extend prior studies by providing a comprehensive framework to analyze the performance limits of wide variety of hyperspectral imaging systems and spectral unmixing algorithms.

